# KIRCLE: An Analysis of Variations in KIR Genes in The Cancer Genome Atlas and UK Biobank

**DOI:** 10.1101/2021.09.02.458787

**Authors:** Galen F. Gao, Dajiang Liu, Xiaowei Zhan, Bo Li

**Author notes:** denotes corresponding author.

## Abstract

**Background:** Natural killer (NK) cells represent a critical component of the innate immune system’s response against cancer and viral infections, among other diseases. To distinguish healthy host cells from infected or tumor cells, killer immunoglobulin receptors (KIR) on NK cells bind and recognize Human Leukocyte Antigen (HLA) complexes on their target cells. Just like the HLAs they bind, these KIRs exhibit high allelic diversity in the human population.

**Results:** In order to better understand these immunoreceptors, we have developed KIRCLE, a novel method for genotyping individual KIR genes from whole exome sequencing data, and used it to analyze approximately 60,000 patient samples in The Cancer Genome Atlas and UK Biobank. We were able to assess population frequencies for different KIR alleles and demonstrate that, similar to HLA alleles, individuals’ KIR alleles correlate strongly with their ethnicities. In addition, we observed associations between different KIR alleles and HLA alleles, including HLA-B*53 with KIR3DL2*013 (Fisher’s Exact FDR = 7.64e-51). Finally, we showcased statistically significant associations between KIR alleles and various clinical correlates, including peptic ulcer disease (Fisher’s Exact FDR = 0.0429) and age of onset of atopy and various KIR alleles (Mann-Whitney-U FDR = 0.0751).

**Conclusions:** KIR polymorphism and NK cells play a critical role in many diseases, often through their interactions with HLA complexes. Peptic ulcer disease and atopy are just two diseases in which NK cells may play a role beyond their “classical” realm of anti-tumor and anti-viral responses.

## 1 Background

Natural killer (NK) cells are an important component of the innate immune system that classically play an important role in the body’s anti-tumor and anti-viral responses. In addition to their functions in these processes, recent research has further implicated their involvement in a much wider range of pathological processes that include cardiac, metabolic, oral, and gastrointestinal diseases. [1-4] While they represent only a small minority of circulating lymphocytes (10-15%), NK cells nonetheless are considered to be the immune cell subtype most effective at monitoring and clearing diseased cells from the body.[5]

As one mechanism to distinguish healthy host cells from infected or tumor cells, NK cells employ killer immunoglobulin receptor (KIR) proteins on their membrane surfaces to bind to and recognize Human Leukocyte Antigen Class I (HLA-I) complexes on the surface of their target cells. 15 KIR genes and 2 KIR pseudogenes have been discovered.[6] These 15 genes may broadly be categorized into either activating KIRs, which promote NK cell activation and induce killing of the target cell on receptor stimulation, or inhibitory KIRs, which prevent NK cell activation and spare the target cell upon ligand binding. Inhibitory KIRs generally possess long cytoplasmic tails and are denoted with an L (KIR2DL1, KIR2DL2, KIR2DL3, KIR2DL5A/B, KIR3DL1, KIR3DL2, and KIR3DL3), whereas activating KIRs generally possess short cytoplasmic tails and are denoted with an S (KIR2DS1, KIR2DS2, KIR2DS3, KIR2DS4, KIR2DS5, and KIR3DS1); however, KIR2DL4 uniquely among the 15 KIR genes possesses both activating and inhibitory functions.[7] Modulation of NK cell activity, and thus susceptibility or resistance to various pathologies, likely depends strongly on the binding properties and interactions between KIR and HLA-I molecules. A high level of diversity in NK cell activity and its outcomes may be achieved largely through four different mechanisms: KIR recognition of highly distinct subsets of HLA-I allotypes, combination of KIRs into distinct haplotypes in different individuals, stochasticity of KIR expression on the surface of individual NK cells, and allelic polymorphism of individual KIR genes.[8] In this manuscript we primarily explore the last of these mechanisms and its downstream effects on disease susceptibility by performing KIR allele inference using Next Generation Sequencing (NGS) data.

Previous attempts to perform KIR genotyping at the individual gene level have either 1) relied on specially prepared primers and amplicon design; 2) required manual review as part of the algorithm; 3) utilized an experimental platform completely different from NGS; or 4) have merely assessed KIR gene presence or deletion rather than detected single-nucleotide-variants. [9-12] Given the rise and modern prevalence of NGS, especially with the recent releases of Whole Exome Sequencing (WES) data for large datasets including UK Biobank and The Cancer Genome Atlas (TCGA), there is a strong need for a fully automated pipeline that can detect single-nucleotide variants of these KIR genes using aligned WES data. In this work, we have developed and characterized the performance of a fully automated algorithm for accurate inference of KIR genes alleles from WES data: “KIR CaLling by Exomes” (KIRCLE). To demonstrate the utility of such an automated KIR genotyper, after running KIRCLE on 10,332 TCGA and 49,953 UK biobank exome samples, we discovered several novel correlations between KIR allele calls and other molecular and clinical features in these two datasets. Our work represents the first large-scale genetic analysis to elucidate pathologic and immunologic associations with human natural killer cells and provides an unprecedented resource for future investigations into the functionality of different KIR alleles.

## 2 Implementation

### 2.1 KIRCLE Workflow Description

KIRCLE is an allele inference algorithm that uses aligned WES data in the form of a BAM or CRAM file to generate probability estimates for each KIR allele, as well as genotype predictions for each KIR gene. KIRCLE consists of 4 major steps: pre-processing, local alignment with BLAST, bootstrapped expectation-maximization, and thresholding (**Figure 1a**).

**Figure 1.**
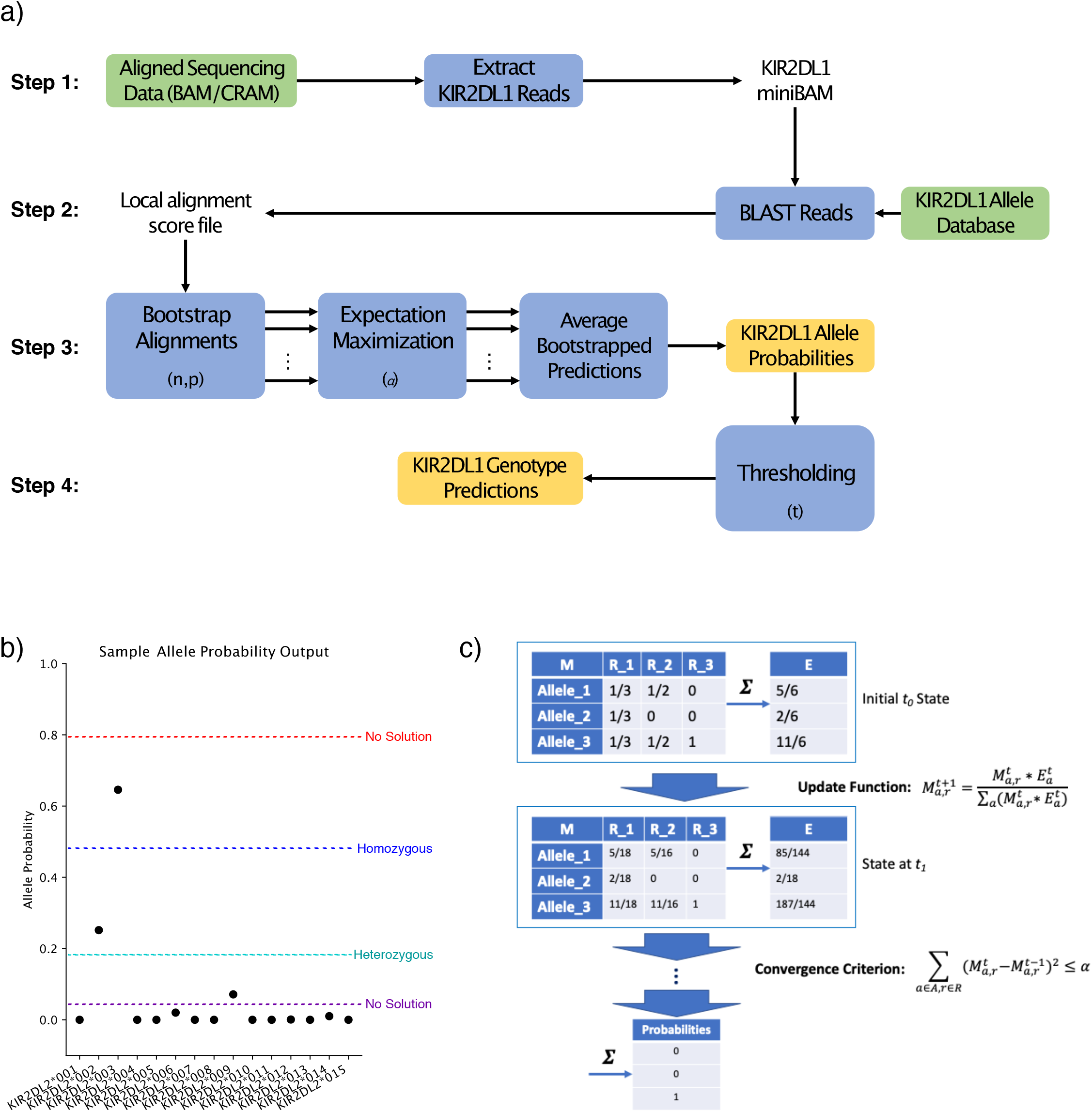
Description of the KIRCLE Methodology. **(a)** Flowchart describing the 4 steps of the KIRCLE algorithm as it processes a single KIR gene (KIR2DL1 as an example here). Inputs are green, computations are blue, and outputs are gold. KIRCLE hyperparameters are listed in parentheses where they are implemented. **(b)** Depiction of step 4 of KIRCLE (thresholding). Allele probabilities generated by expectation-maximization may lead to a homozygous solution, a heterozygous solution, or no solution at all, depending on the selected value of the threshold hyperparameter *t*. **(c)** Depiction of one step of expectation-maximization. The initial allele-read matrix 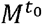 is collapsed into an expectation vector 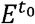 that is used to compute the next iteration of the matrix 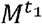. This process is repeated until the convergence criterion is satisfied, at which point the final expectation vector represents an estimate of KIR allele probabilities.

1. In pre-preprocessing, KIRCLE first extracts all WES reads that map to the genomic coordinates of the KIR genes on chromosome 19q13.4 and writes these reads to fifteen separate files—one for each KIR gene.
2. Next, KIRCLE uses nucleotide BLAST to perform local alignment on each KIR gene’s collection of reads against a database of variants belonging to that particular KIR gene. In the IPD-KIR Database v2.8.0, 908 different alleles spanning the 15 KIR genes are documented, of which 535 represent distinct coding variants. KIRCLE then filters out alignments with less than 100% identity matches to documented KIR alleles.
3. KIRCLE then bootstraps the BLAST-identified alignments with 100% identity matches to KIR alleles and uses an expectation-maximization (EM) algorithm, with convergence hyperparameter *α*, to generate allele probability estimates from these collections of alignments. *n* bootstraps of fraction *p* of all 100%-identity alignments are computed in this manner. The bootstrapped allele probability estimates are then averaged together to determine a final probability estimate for each allele. This bootstrapping is helpful in countering the EM algorithm’s tendency to converge to local minima representing homozygous solutions and based on small differences in initial alignment data.
4. Finally, KIRCLE uses a thresholding algorithm to convert each KIR gene’s set of allele probability estimates into homozygous or heterozygous genotype calls, depending on the number of alleles that exceeded a heuristically determined threshold *t* (**Figure 1b**).

Final workflow outputs include a table of allele probability estimates, a table of genotype guesses, and a list of runtime hyperparameters.

### 2.2 Allele Inference Using Expectation Maximization

To infer allele probabilities from a set of read alignments to a database of KIR alleles, we use an expectation-maximization algorithm to aggregate the alignment data into an initial set of allele probability “expectations,” which is then used to further weight the alignment data in order to refine our estimates of KIR allele probabilities. Thus, given a bootstrap of read alignments with 100% identity to at least one KIR allele, KIRCLE’s EM algorithm iteratively generates probability estimates for each KIR reference allele (**Figure 1c**).

Let:

*m* = the total number of alleles of this KIR gene

*n* = the total number of reads in this bootstrap

*A* = the set of KIR alleles {*a*_1,_ *a*_2,_ *a*_*m*_}

*R* = the set of BAM reads {*r*_1,_ *r*_2,_ *r*_*n*_}

*x*_*r*_ = the number of alleles that read *r* aligns to with 100% identity

*t* = each time step of the expectation maximization algorithm

*α* = a heuristically chosen convergence threshold

We first initialize an *m* x *n* “alignment matrix” *M* to encode our read alignments:

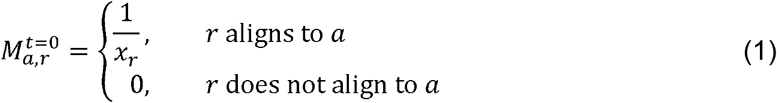

Next, using *M*, we compute an initial expectation vector *E*^*t*^ representing our rudimentary estimate of each KIR allele’s probability in this sample:

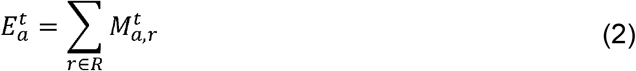

Then, at each time step *t* of the expectation-maximization algorithm, we update the values of our alignment matrix *M* in a Bayesian fashion using the previously generated expectation vector *E*^*t*^ as our prior and *M*^*t*^ as our likelihood:

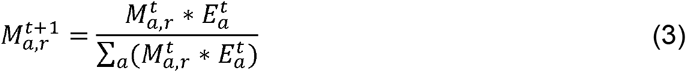

Subsequently, we may generate an updated expectation vector *E*^*t+1*^ using *M*^*t+1*^ in conjunction with equation (2) above.

We continue to iterate through our expectation-maximization algorithm in this manner, computing *E*^*t*^ from *M*^*t*^ and then *M*^*t+1*^ from *E*^*t*^ and *M*^*t*^, until we achieve our convergence criterion, defined as the sum of squared changes in *M* not exceeding a heuristically selected hyperparameter *α*:

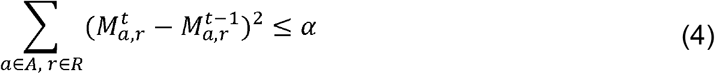

Ultimately, our final expectation vector *E*^*T*^ is outputted as our vector of allele probability estimates.

### 2.3 Allele Coding Region Collapse

Because we used WES data as our input, KIR variants that differed only at non-exonic sites were merged by summing their allele probability estimates. Furthermore, as we are primarily interested in the phenotypic effects of altered binding affinity to KIR domains, we merged variants that differ only by a silent mutation by summing their allele probability estimates as well. Thus, all KIR alleles subsequently are reported as a three-digit number following the KIR gene name (e.g. KIR2DL4*005).

### 2.4 Clinical Effect Model Comparison

To explore several correlations we discovered between KIR3DL2 genotype and earliest age of atopy onset more deeply, we attempted to model earliest age of atopy onset as a linear function of the KIR3DL2 genotype

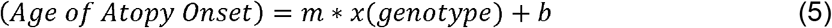

where *x(genotype)* is determined by the model of KIR allele expression. Under a dominant model, both homozygotes and heterozygotes for an allele KIR3DL2*000 contribute equally to the phenotype. Thus,

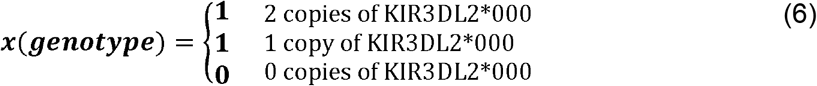

Meanwhile, under a semi-dominant model, homozygotes are twice as expressive as heterozygotes:

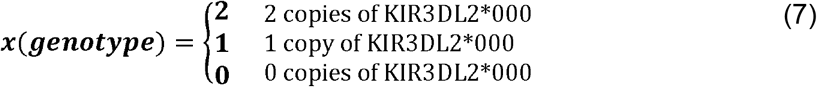

Finally, under a recessive model, only homozygotes have an effect on phenotypic expression:

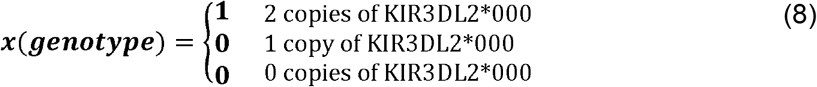

How well each of these models performed against each other was assessed by the goodness of fit of the final linear model (equation 5) with the UK Biobank data and summarized using the Bayesian Information Criterion.

## 3 Results

### 3.1 Hyperparameter Determination and Validation for KIRCLE

KIRCLE requires the use of 4 hyperparameters: *α* (the convergence threshold for expectation-maximization), *n* (the number of bootstraps to perform), *p* (the proportion of reads to use in each bootstrap) and *t* (the threshold used to convert KIR allele probabilities to binary KIR genotype calls). Of these, choices regarding *p* and *t* represent the greatest and most direct potential sources of variability in KIRCLE’s accuracy. Using one randomly selected sample from UK Biobank, we were able to characterize KIRCLE’s performance, as measured by the Shannon entropy of the inferred genotypes, across different values of *p* (from 0.2 to 0.8) and *t* (from 0.05 to 0.40). At each set of hyperparameters tested, we performed 500 iterations of KIRCLE on one arbitrarily selected sample in UK Biobank, collected the 500 genotype outputs, and empirically computed the log2 Shannon entropy of the genotype solutions for each KIR gene. An ideal genotype caller would be consistent and call the same solution for the same input, resulting in a “genotype-entropy” of 0. For many KIR genes, such as KIR2DL1 and KIR2DL4, contour maps of the resulting entropies revealed that KIRCLE was largely self-consistent, with little variability of output (genotype-entropy of 0) across a wide spectrum of hyperparameter values (**Figure 2a-b**). This pattern was recapitulated in the majority of KIR genes, suggesting respectable consistency of KIRCLE output across multiple KIR genes (**Supplementary Figure S1a-j**). For all subsequent analyses in this manuscript, hyperparameter values of *α*=1e-5, *n*=100, *p*=0.5, and *t*=0.25 were used.

**Figure 2.**
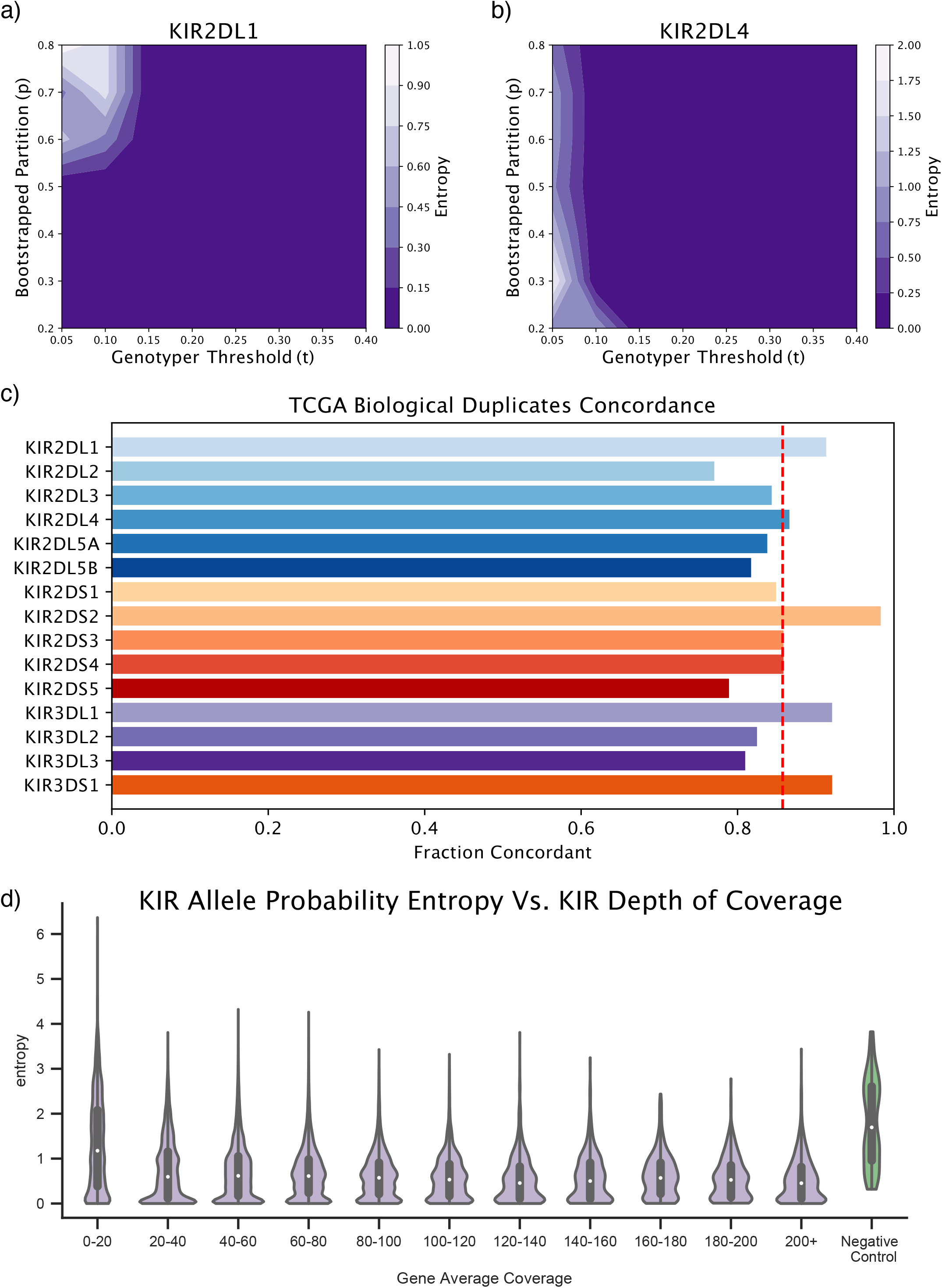
KIRCLE Accuracy and Consistency Validation. **(a)** Contour plot demonstrating the effect of varying the bootstrap-proportion (*p*) and threshold (*t*) hyperparameters on KIR2DL1 allele inference, as measured by empirical calculation of the inferred genotypes’ entropy. **(b)** KIRCLE’s performance on KIR2DL4 allele inference was similarly characterized. **(c)** Fraction of each KIR gene’s KIRCLE-inferred allele genotypes that were called identically between 531 samples and their biological replicates in TCGA. **(d)** TCGA sample coverages (binned) versus TCGA sample allele probability entropies for all 15 KIR genes. The allele probability entropies of a set of 20 “pseudo-BAMs” (green) are presented as negative controls.

Next, to establish the accuracy of our algorithm, we assessed the concordance of KIRCLE-generated genotypes between TCGA biological replicates. Of 10,332 exomes in TCGA, 1,062 were present twice as biological replicates and thus used in this analysis. We determined that 85.8% of genotype solutions called by KIRCLE across all KIR genes were concordant between replicates (**Supplementary Figure S2a**). We defined solutions to be concordant if the genotype inferred by KIRCLE in one sample was identical to that inferred in its replicate. Genes with the highest concordance between replicates were KIR2DS2 (98.3%), KIR3DL1 (92.1%), and KIR3DS1 (92.1%), whereas genes with the lowest concordance between replicates were KIR2DL2 (77.0%), KIR2DS5 (78.9%), and KIR3DL3 (81.0%) (**Figure 2c**).

Finally, we investigated whether KIRCLE is robust against differences in depth of sequencing. To do so, we compared the ambiguity of KIRCLE’s output, quantified as the Shannon entropy of generated KIR allele probabilities, across TCGA samples with different depths of sequencing. Low ambiguity in KIR allele calling results in KIR allele probabilities of either 1 for a single allele and 0 for all other alleles (reflecting a homozygous genotype) or 0.5 for two different alleles and 0 for all other alleles (reflecting a heterozygous genotype), leading to entropies of either 0 or 1 respectively. Conversely, high ambiguity in KIR allele calling will lead to a more uniform distribution of KIR allele probabilities, leading to entropies higher than 1. For each TCGA sample, we measured both the average coverage and the KIR-allele-probability entropies for each KIR gene. Binning samples by their average coverages, we observed that allele probability entropies—and thus the ambiguity of KIR allele probabilities—are notably increased only at very low coverages (<20 average depth of coverage at the KIR gene locus). Furthermore, as negative controls, 20 “pseudo-BAMs” were generated by randomly sampling reads mapping to KIR gene loci from 50 randomly selected BAMs in TCGA. Pseudo-BAMs were generated with an average read depth commensurate with their constituent BAMs. After applying KIRCLE to these pseudo-BAMs, their resulting allele probability entropies were much higher (median=1.70) than a significant majority of actually observed entropies for all TCGA samples, regardless of the depth of coverage (**Figure 2d**). Moreover, despite differences in average depth of coverage at different KIR gene loci, average KIR allele entropies between different KIR genes largely remained constant (**Supplementary Figure S2b**). Overall, KIRCLE demonstrated a high level of consistency while being able to call a diverse set of KIR genotype solutions and is robust to the effects of low depth of coverage.

### 3.2 KIR Allele Comparisons Between TCGA and UK Biobank

After benchmarking KIRCLE using internal quality control metrics, we assessed KIRCLE’s performance by comparing its allele predictions in TCGA to its allele predictions in UK Biobank. We first compared the frequencies of different KIR alleles in TCGA with their frequencies in UK Biobank. For each KIR gene, we ranked its alleles by frequency in both TCGA and UK Biobank and then computed the Spearman correlation coefficient between the allele frequencies in the two datasets (**Figure 3a**). We noted that all KIR genes displayed positive correlation coefficients and that the vast majority of KIR genes demonstrated highly similar distributions of allele frequencies between TCGA and UK Biobank (Spearman’s ρ=0.800). Direct comparison of all KIR alleles ranked by frequency also demonstrated high consistency between the two cohorts (**Figure 3b**). Although both UK Biobank and TCGA are largely composed of Caucasians (81.4% and 93.3% of the individuals we analyzed in TCGA and UK Biobank respectively), there exist small yet notable differences in ethnic makeup between the two datasets. These differences in ethnic compositions may account for a subset of the observed differences in KIR allele frequency.

**Figure 3.**
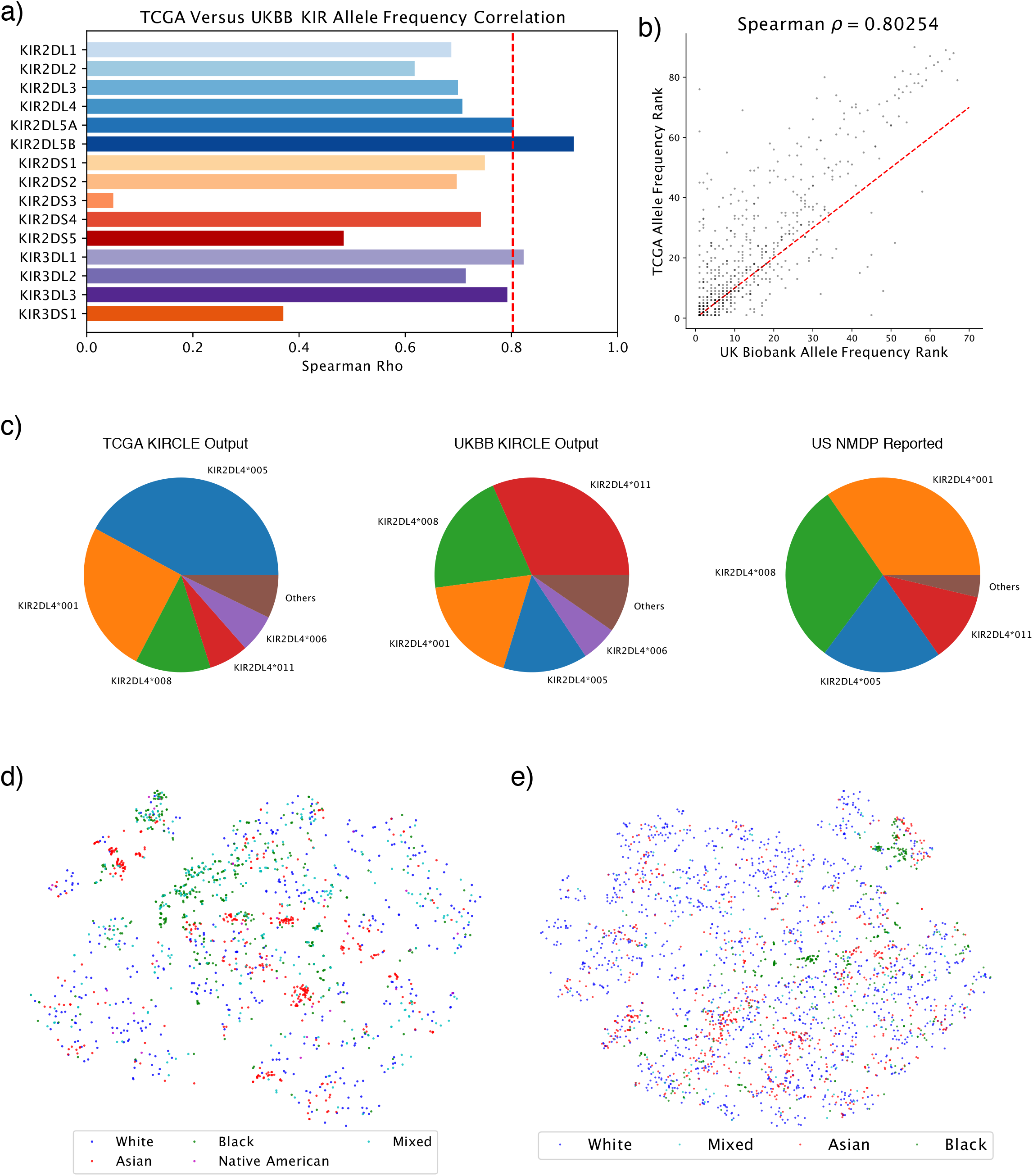
KIR Allele Distributions in UK Biobank and TCGA. **(a)** Non-linear correlations between KIR allele frequencies in TCGA versus those in UK Biobank, stratified by KIR gene. **(b)** Comparison of allele frequency ranks between TCGA and UK Biobank. **(c)** KIR2DL4 allele frequencies in TCGA (left), UK Biobank (center), and US NMDP (right). **(d)** t-SNE plot of individuals in TCGA colored by participants’ ethnicities. Caucasian individuals were down-sampled by a factor of 16 for ease of visualization. **(e)** t-SNE plot of UK Biobank individuals colored by ethnicity. Caucasian individuals again were down-sampled by a factor of 16 for ease of visualization.

Additionally, we were able to further validate observed KIR allele frequencies for certain KIR genes using allele frequency data from the United States National Marrow Donor Program (NMDP), as reported by the Allele Frequency Net Database.[13] We used the NMDP dataset because the subjects in this cohort are predominantly Caucasians, similar to the TCGA patients. For KIR2DL4, the four most frequent KIR2DL4 alleles reported by the NMDP were KIR2DL4*001, KIR2DL4*008, KIR2DL4*005, and KIR2DL4*011 (34.6%, 30.2%, 19.9%, and 11.6% respectively). We were able to recapitulate these four alleles as the most frequent KIR2DL4 alleles in both TCGA and UK Biobank, albeit in a different order for each dataset (**Figure 3c**). In TCGA, KIR2DL4*005 was the most frequent allele, followed by KIR2DL4*001, KIR2DL4*008, and KIR2DL4*011 (42.0%, 25.4%, 12.4%, and 6.58%). In UK Biobank, this order was reversed with KIR2DL4*011 being the most frequent allele, followed by KIR2DL4*008, KIR2DL4*001, and finally KIR2DL4*005 (31.6%, 20.5%, 18.2%, and 14.0%). Further validation of allele frequencies against the NMDP was also performed for the alleles of KIR3DL2. The most frequent KIR3DL2 allele in a population of 75 Caucasians was KIR3DL2*002 (26.1%), followed by KIR3DL2*001 and KIR3DL2*007 (21.0% and 18.8%).[14] While KIR3DL2*002 was found at similarly high frequencies in both TCGA (9.85%) and UK Biobank (9.27%) as the 2^nd^ and 3^rd^ most frequent KIR3DL2 alleles respectively, KIR3DL2*001 and KIR3DL2*007 were much lower ranked at 8^th^ and 9^th^ in TCGA and 8^th^ and 1st in UK Biobank respectively. However, these are still fairly well-represented alleles at 4.94% and 2.67% frequency in TCGA and 5.12% and 13.3% frequency in UK biobank respectively. Furthermore, considered overall, KIR3DL2 allele frequency ranks in TCGA and UK Biobank still demonstrate positive correlations with the allele frequency ranks observed in the NMDP (**Supplementary Figure S2c**). Despite slight numerical differences, confirmation of the status of the most frequent alleles in these two KIR genes increases our confidence in KIRCLE’s ability to infer KIR alleles from WES data accurately.

In addition to validating population frequencies of KIR alleles, we also examined patterns of KIR allele co-expression and dependence. As KIRCLE assesses for the presence of 535 KIR alleles over 15 KIR genes, the KIR genotype of each sample in TCGA and UK Biobank may be represented as a point in 535-dimensional “KIR-space.” We first used t-distributed stochastic neighbor embedding (t-SNE) to perform dimensionality reduction and thus visualize the distribution of individuals in TCGA in 2 dimensions.[15] When we colored this t-SNE map using individuals’ SNP-inferred ethnicities,[16] we observed that different ethnicities cluster together and are non-uniformly distributed (**Figure 3d**). In particular, African Americans and—to a lesser extent—Asian Americans in TCGA formed clusters that were often very distinct from the Caucasian majority. Similar analyses performed in UK Biobank recapitulated this non-random distribution of KIR genotypes and confirmed the non-uniform distribution and clustering of those who self-identified their ethnicity as “Black” or “Asian” (**Figure 3e**). Of particular note, the “Asian” population in TCGA comprises those of East Asian descent, whereas the “Asian” population in UK Biobank largely comprises those of South Asian descent (with major subcategories of Indians, Pakistanis, and Bangladeshis). However, both groups of Asians clustered distinctly and separately from the Caucasian majority to some extent in both datasets.

### 3.3 KIR Allele Associations with HLA Alleles

As it is known that HLA and KIR bind to each other in an allele-specific way, we posited that strong correlations may also exist between KIR alleles and HLA alleles on the population level, due to a known co-evolution event in humans.[17] Using HLA types imputed by the HLA*IMP:02 algorithm and subsequently released by UK Biobank,[18] we observed 326 significantly associated pairs of KIR alleles with HLA alleles in the UK Biobank data (**Figure 4a-b, Supplementary Figure S3a**). Many of these associations belonged to a set of particularly common HLA alleles (e.g. HLA-B*53) or KIR alleles (e.g. KIR3DL3*005). Furthermore, we also note that the majority (73.0%) of significant associations are positive. We speculate that these associations reflect changes in direct physical interactions between HLA and KIR alleles, which result in co-selection due to an advantageous increase in fitness for individuals with these combinations of KIR and HLA alleles. Particularly visually striking examples of positive and negative associations between KIRs and HLAs include: KIR3DL3*005 with HLA-A*74 (Fisher’s Exact FDR=7.43e-43; odds ratio=55.1) and KIR2DL3*002 with HLA-A*36 (Fisher’s Exact FDR=1.71e-12; odds ratio=0.0727) respectively (**Figure 4c**). Additionally, when examining the t-SNE coordinates of individuals with HLA alleles such as HLA-B*42, we observed a non-uniform distribution and clustering of these samples that closely mirrors the distribution of samples when labeled by ethnicity (**Figure 4d**).

**Figure 4.**
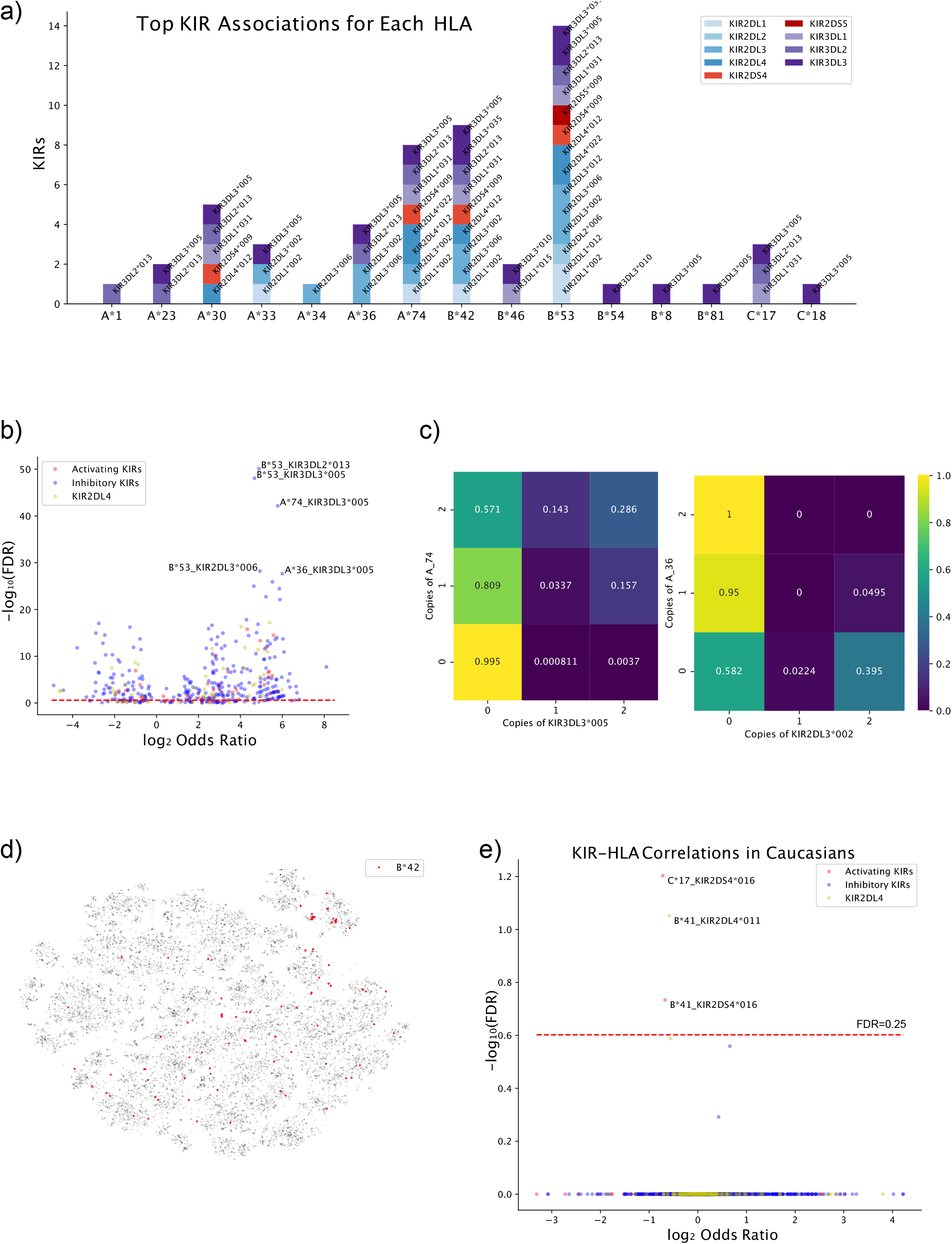
KIR Allele Associations with HLA Alleles. **(a)** Bar plot depicting and listing KIR alleles that significantly associated with HLA alleles at the FDR < 1e-10 level. **(b)** Volcano plot of KIR allele correlations with HLA alleles in UK Biobank. Associations are color-coded by the activity (inhibitory or activating) of the KIR allele. **(c)** Presence of at least 1 copy of HLA-A74 positively correlates with presence of at least 1 copy of KIR3DL3*005 (left) and presence of at least 1 copy of HLA-A36 negatively correlates with presence of at least 1 copy of KIR2DL3*002. **(d)** KIR t-SNE of UK Biobank individuals with those possessing HLA-B*42 highlighted in red. **(e)** Volcano plot of KIR allele correlations with HLA alleles among Caucasians in UK Biobank.

While these findings may support the biological link between these two classes of molecules and shed additional light onto which particular HLA alleles may have evolved in parallel with particular KIR alleles, they also raise the possibility that our observed associations are driven by population stratification according to ethnicity. In order to disentangle the effects of this stratification on associations between HLA and KIR alleles, we re-attempted the analysis using only Caucasian individuals in UK Biobank, while testing only KIR alleles with >1% allele frequency in UK Biobank (**Figure 4e**). While this analysis unveiled a much smaller subset of HLA-KIR associations, we noted 3 significant associations: HLA-C*17 with KIR2DS4*016 and HLA-B*41 with KIR2DL4*011 and KIR2DS4*016. Notably, both KIR2DS4 and KIR2DL4 have NK-cell-activating activity, and all three are affiliated with a negative odds ratio. These results indicate that HLA-C*17 and B*41 could be true activation ligands for KIR2DS4 and KIR2DL4, and their interactions may induce NK responses that impose negative selection pressure on individuals bearing both alleles.

Although TCGA is a much smaller dataset than UK Biobank, we were able to use TCGA to discover a smaller set of correlations between HLA alleles and KIR alleles after filtering out KIR alleles with <1% allele frequency in TCGA to improve our Bonferroni correction factor (**Supplementary Figure S3b**). HLA allele calls for samples in TCGA were made using POLYSOLVER.[19] In particular, KIR2DL2*003, KIR3DL2*013, and KIR3DL3*008 were strongly positively associated with HLA-B*46, HLA-B*53, and HLA-C*15 respectively at the FDR < 0.25 level. The HLA-B*53 association with KIR3DL2*013, notably, was the most significant HLA-KIR association discovered in UK Biobank. However, when we re-attempted the analysis using only Caucasian individuals in TCGA to eliminate population stratification by ethnicity as a potential confounding factor, all significant associations between KIR and HLA alleles disappeared after Bonferroni correction. In summary, after correction of population stratifications, we found few significant associations between activating KIR gene and HLA alleles. The absence of significant associations between inhibitory KIR genes and HLA alleles might suggest weaker selective pressure for KIR alleles, possibly due to the multiple redundant mechanisms inhibiting NK cell activation. [20]

### 3.4 KIR Allele Associations with Clinical Correlates

In addition to correlations with HLA alleles, we searched for KIR allele correlations with clinical features. We first examined KIR allele correlations with individuals’ medical diagnoses documented in UK Biobank, as encoded by the 10^th^ revision of the International Statistical Classification of Diseases (ICD10). To minimize the number of under-powered tests we performed, we attempted correlations only with KIR alleles represented at over 1% frequency in UK Biobank. Additionally, we excluded all diseases primarily associated with external causes, including accidents, injuries, and nutritional deficiencies, as well as obstetric and psychiatric diseases among others. Of note, this list of exclusions includes infectious diseases, which despite having a strong biological basis for association with KIR alleles, require exposure to a pathogen, which is largely driven by individuals’ environmental circumstances. Strikingly, the only associations that remained significant at the FDR < 0.25 level were those associated with sickle-cell anemia (ICD10 D57) or with uterine leiomyomas (ICD10 D25), both diseases that disproportionately affect black people.[21] However, positing a direct biological mechanism behind these associations likely would represent a third-cause fallacy, as blacks are statistically more likely to possess both KIR alleles enriched in black populations as well as either the sickle-cell trait or uterine leiomyomas.

Thus, we next narrowed our analysis to investigate only those individuals who self-identified as Caucasian. While the vast majority of correlations failed false-discovery-rate correction, we discovered a significant correlation between the KIR3DL3*080 allele and ICD10 K25—Peptic Ulcer Disease (PUD) (Fisher’s Exact test, FDR= 0.0429; **Figure 5a**). Whereas those without KIR3DL3*080 had merely a 1.04% chance of being diagnosed with PUD, patients with KIR3DL3*080 had a 2.90% chance of being diagnosed with PUD, representing a 2.8-fold increase in likelihood (**Figure 5b**). No significant association was found between KIR3DL3*080 and usage of ibuprofen, which would predispose individuals toward developing PUD (data not shown). Thus, if KIR3DL3*080 predisposes an individual toward PUD, it likely does so through an alternative mechanism. Significant correlations with ICD10 diagnosis codes in Black and Asian populations were not observed, likely owing to the lower statistical power these smaller populations had.

**Figure 5.**
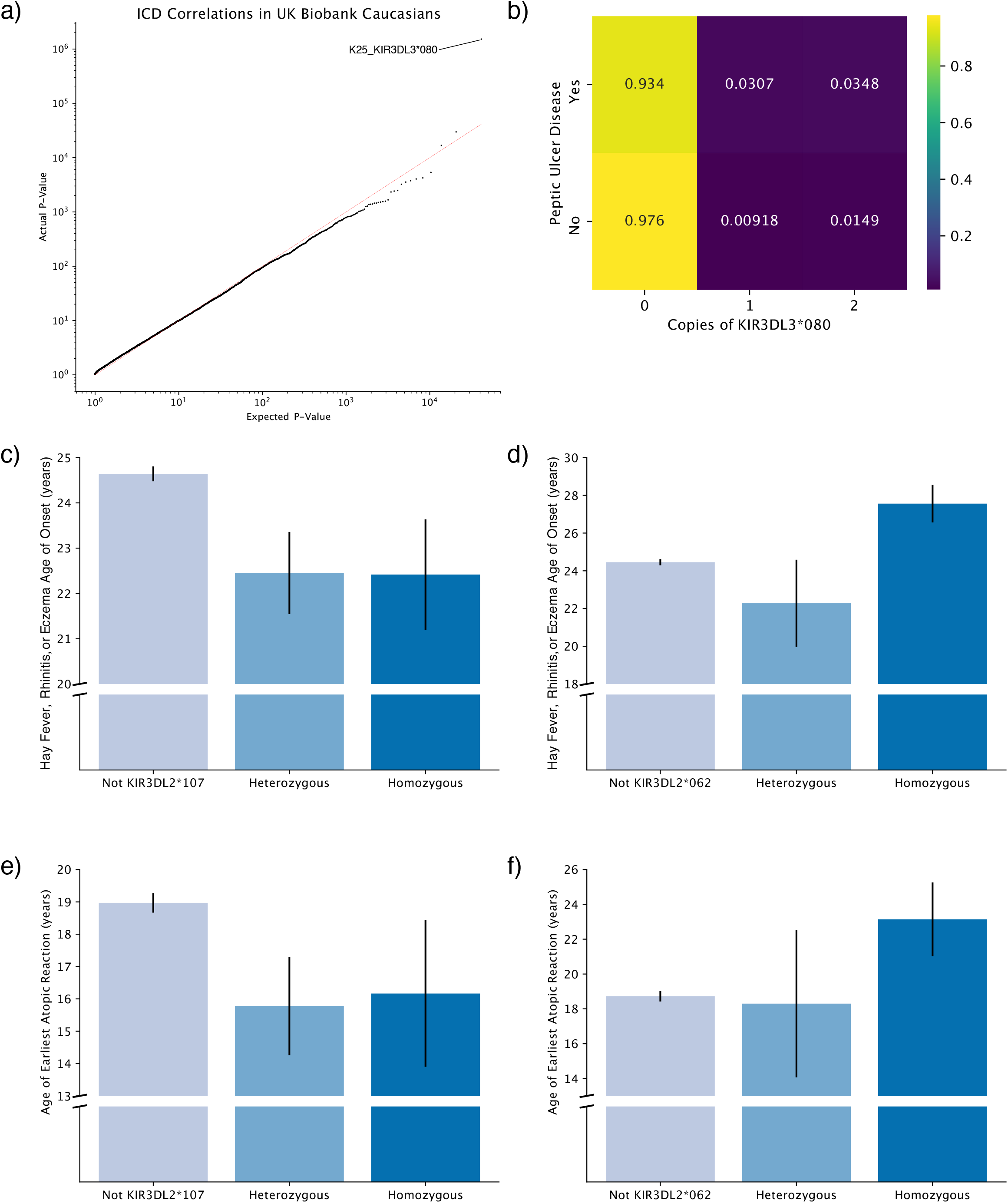
KIR Allele Associations with Clinical Correlates. **(a)** QQ-plot of KIR allele correlations with ICD10 diagnosis codes in UK Biobank among Caucasian individuals. **(b)** Odds of developing peptic ulcer disease is increased among those with the KIR3DL3*080 phenotype. **(c)** Bar plot showing decreased mean age of hay fever, rhinitis, or eczema in those with at least one copy of KIR3DL2*107. **(d)** Bar plot showing increased mean age of hay fever, rhinitis, or eczema in those homozygous for KIR3DL2*062. **(e)** Bar plot showing decreased mean age of atopy in those with at least one copy of KIR3DL2*107. **(f)** Bar plot showing increased mean age of atopy in those homozygous for KIR3DL2*062.

Additionally, we explored correlations between KIR alleles with population frequency >1% and other clinical correlates besides ICD10 codes. When examining correlations with age of onset of several chronic diseases and conditions, we discovered that KIR3DL2*107 was highly correlated with early age of onset of hay fever, rhinitis, or eczema in Caucasian individuals. Whereas individuals without KIR3DL2*107 had an average age of onset of 24.7 years, those with at least one copy of KIR3DL2*107 had an average age of onset of 22.4 years (two-sided Mann-Whitney test, FDR=0.0751; **Figure 5c**). Moreover, an alternative allele of KIR3DL2, KIR3DL2*062, was weakly associated with an increase in age of onset of hay fever, rhinitis, or eczema from 24.5 years to 27.0 years (Mann-Whitney FDR=0.244; **Figure 5d**). Later onset of these conditions was particularly pronounced in individuals with two copies of KIR3DL2*062 (average age of onset of 27.6 years). Together, these results suggest that polymorphisms in KIR3DL2 may play a key role in determining the age of onset of hay fever, rhinitis, and/or eczema.

Moreover, hay fever and eczema, in conjunction with allergic asthma, more broadly represent manifestations of atopy, the genetic predilection to trigger IgE-mediated (Type I) hypersensitivity reactions following allergen exposure with increased T_H_2-driven responses.[22] Thus, we next attempted to generalize this association to encompass atopy more broadly by examining KIR3DL2*107 and KIR3DL2*062’s associations with age of onset of either asthma or hay fever, rhinitis, or eczema, using the age of onset of whichever condition occurred earliest in life for each individual. We observed the same association: individuals with at least one copy of KIR3DL2*107 had an average age of onset of 15.9 years, whereas those without any copies of KIR3DL2*107 had an average age of onset of 19.0 years (Mann-Whitney p=0.012; **Figure 5e**). Simultaneously, individuals with at least one copy of KIR3DL2*062 (22.5 years), and particularly those with two copies of KIR3DL2*062 (23.1 years), had later onsets of atopic reactions than those without KIR3DL2*062 (18.7 years; Mann-Whitney p=0.019; **Figure 5f**). Together, these findings suggest a potential biological mechanism either delaying or hastening onset of atopic reactions like hay fever, eczema, or asthma that involves KIR3DL2, and the KIR3DL2*107 and KIR3DL2*062 alleles in particular. In addition to atopic reactions, we also observed significant associations of KIR alleles with other clinical correlates, including dental and oral health, quantitative blood analysis, and waist circumference, suggesting potentially broad impact of natural killer functions in affecting diverse human traits (**Supplementary Figure 4**).

Finally, to further explore the effects of KIR3DL2 polymorphism on age of atopy onset, we posited that each of the two aforementioned KIR3DL2 alleles follows either a dominant, semi-dominant, or recessive model of expression and then sought to determine which of these three models best explains the effect of KIR3DL2 genotype on age of atopy onset. In the recessive model, only a genotype homozygous for the KIR3DL2 allele in question contributes to a change in age of atopy onset from baseline. In contrast, in the dominant model, genotypes either homozygous or heterozygous for the KIR3DL2 allele in question contribute to changes in baseline age of atopy onset. Finally, in the semi-dominant model, homozygotes for the KIR3DL2 allele in question are twice as potent as corresponding heterozygotes in changing age of atopy onset from baseline. When assessed against each other using the UK Biobank data, the dominant model outperformed semi-dominant and recessive models of expression for KIR3DL2*107, as measured by the Bayesian information criterion (−10.8145 versus -10.8151 and -10.8172 respectively). Meanwhile, expression patterns of KIR3DL2*062 instead favored the recessive model over the semi-dominant and dominant models of expression for KIR3DL2*062, as measured by the Bayesian information criterion (−10.8138 versus -10.8141 and -10.8146 respectively). In summary, our analysis indicated that KIR3DL2*107 may “override” other alleles and thus present with a dominant phenotype, whereas KIR3DL2*062 may be weaker than other KIR3DL2 alleles and thus present with a recessive phenotype.

## 4 Discussion

The fifteen KIR genes represent a polymorphic set of immune modulators with an array of potential effects on immune and clinical phenotypes. In this manuscript, we have developed, characterized, and implemented our algorithm KIRCLE, uncovering multiple correlations between KIR alleles and other features in TCGA and UK Biobank.

### 4.1 KIR Alleles Associated with HLA Alleles

Class I HLAs represent well-known binding partners of KIRs. Thus, any change in either the KIR binding site or the HLA binding site that alters their affinities to each other may be expected to modulate NK cell activation or inhibition. However, as both HLA and KIR loci are highly polymorphic, it has historically been challenging to determine their matches through low-throughput experimental approaches or through small-scale computational analyses. By using KIRCLE and a large cohort of UK Biobank data, we were able to observe statistically significant interacting KIR and HLA allele pairs, which were previously unknown. Specifically, we observed many KIR alleles to be more frequently expressed in individuals who also possess particular HLA alleles, which might be indicative of selective pressures for these receptors to co-evolve to maintain appropriate levels of NK cell activity.

However, in addition to observing differences in KIR allele frequencies among those with different HLA alleles, we also observed differences in KIR allele frequencies among different ethnic populations, which are already known to possess different HLA allele frequencies.[23] While this combination of observations may reflect common selective pressures that were historically experienced by these ethnic populations which then may have forced KIR and HLA alleles to co-evolve, they also pointed to ethnic stratification as a potential confounding factor in our purely correlative study. However, when we removed this potential confounder by examining only Caucasian individuals in UK Biobank, we observed that HLA-C*17 and HLA-B*41 are found at lower frequency in individuals with KIR2DS4*016 and KIR2DL4*011, two KIR alleles with known activating activity, than in those without. We posit that this “anti-correlation” may represent evidence of an intolerance or aversion toward potentially lethal NK cell hyperactivity. Thus, individuals with these particular KIR-HLA allele combinations may be underrepresented due to over-activation of NK cells.

### 4.2 KIR Allele Associations with Peptic Ulcer Disease and Atopic Reactions

Previous genome-wide association studies of PUD have largely been performed in East Asian populations and did not uncover any associations between KIR polymorphism and either PUD or *H. pylori* infection.[24, 25] However, NK cells are known to be present in the gastric and duodenal mucosa and have been shown to be directly activated by *H. pylori* bacteria to produce IFN-*γ* and trigger an immune response.[26] Our result builds upon these existing known interactions and suggests that the KIR3DL3*080 may increase susceptibility to PUD through modulating NK cells’ natural response to *H. pylori*.

Furthermore, we uncovered a potential association between age of presentation of atopy and two different variants of KIR3DL2: KIR3DL2*107 and KIR3DL2*062. At least one copy of one of these two variants is present in 7.89% of the Caucasian population in UK Biobank. Indeed evidence exists for NK cells’ involvement in atopic and autoimmune diseases of the skin (i.e. eczema), even if the details of this involvement remain unclear, [27] and increasing support has been seen for their role in allergen-specific immune suppression, Th1 cell generation, and Ig production. [28] Our finding that KIR3DL2*107 hastens presentation of atopy and that KIR3DL2*062 delays it potentially further points to a role specifically for KIR3DL2 in regulating NK cell activity as it contributes to these diseases. One possible explanation for the opposite directions of impact on age of onset is that KIR3DL2*107 is stronger than other alleles and thus presents with a dominant phenotype, whereas KIR3DL2*062 is weaker than others and thus results in a recessive phenotype. As preliminary evidence supporting this explanation, we demonstrated that a dominant model of expression best fits KIR3DL2*107, while a recessive model of expression best fits KIR3DL2*062. Nonetheless, future cohorts with larger sample sizes will be required to test this hypothesis further.

### 4.3 Limitations and Future Directions

While we were able to use biological replicates in TCGA to benchmark the accuracy of KIRCLE and compare our population-level estimates of KIR allele frequencies to prior estimates of KIR allele frequencies as reported by the US NMDP to validate our KIR genotype predictions, we were unable to carry out any experimental validation to benchmark its accuracy more directly. Additionally, after generating KIR genotype predictions, our downstream correlations all represented univariate analyses, due to the relatively low abundance of individual KIR alleles. While such a simplistic analysis is suitable for a first-pass search for potential direct associations, more nuanced future analyses of clinical associations with KIR alleles will need to account for confounding factors beyond human genetics to determine individuals’ susceptibility to diseases, including individuals’ living or occupational environments, full medical histories, lifestyles, and much more. Such analyses will likely become available with sufficient statistical power when UK Biobank fulfills its mission to sequence all 500,000 individuals. Furthermore, Caucasian individuals are heavily over-represented in both the TCGA and UK Biobank cohorts, and thus our downstream analyses have largely been suitably powered to investigate only those KIR alleles that are well represented among Caucasians. Future studies will be needed to use more racially diverse cohorts to analyze KIR alleles that are more frequently represented in other ethnicities. Finally, as mentioned at the outset, NK cell activity is modulated by a number of factors outside of individual KIR genes polymorphism, including the subset of HLA-Is they recognize, the distinct combinations of genes that constitute the individual’s KIR haplotype, and stochasticity in KIR expression on the surface of individual NK cells. Indeed, over 40 distinct KIR haplotypes, each composed of at least seven KIR genes, have been documented in the human population.[29] Variation of any of these additional factors may further affect NK cell function and ideally would be explored in conjunction with KIR polymorphism at the individual gene level in future studies.

In conclusion, our work has generated KIR allele predictions for TCGA and UK Biobank that will be invaluable for future studies of NK cells in these populations, uncovered multiple novel associations between KIR gene variants and clinical and molecular features, and has paved the way for future investigation into the role of KIRs in immunologic response and human disease. We hope that our algorithm can serve as a benchmark for future algorithms that will perform KIR genotyping, and that others may use our algorithm to better understand the immunologic and pathologic processes surrounding KIR genes.

## 5 Conclusions

We have developed KIRCLE, a first-of-its-kind fully automated computational pipeline for the inference of germline variants of the highly polymorphic killer-cell immunoglobulin-like receptor (KIR) genes from whole exome sequencing data. We demonstrate the utility of such an algorithm by using KIRCLE to infer germline KIR variants in approximately sixty-thousand individuals in The Cancer Genome Atlas and UK Biobank and then discover novel molecular and clinical correlations with these variants. This work represents the first large-scale genetic analysis to elucidate immunologic and pathologic associations with human natural killer cells and will serve as a valuable resource for future investigations into the immunogenomics and disease processes involving KIRs.

## 6 Availability and Requirements

Project name: KIRCLE

Project home page: https://github.com/gaog94/KIRCLE

Operating system(s): Platform independent

Programming language: Python3

Other requirements: Anaconda, samtools 1.3.1, and Nucleotide BLAST v2.8.1+

License: MIT

Any restrictions to use by non-academics: None.

### 7 List of Abbreviations

NK: Natural Killer
KIR: Killer Immunoglobulin Receptor
HLA: Human Leukocyte Antigen
NGS: Next Generation Sequencing
KIRCLE: KIR CaLling by Exomes
WES: Whole Exome Sequencing
TCGA: The Cancer Genome Atlas
NMDP: National Marrow Donor Program
FDR: False Discovery Rate
t-SNE: t-distributed Stochastic Neighbor Embedding
ICD10: International Statistical Classification of Diseases 10
PUD: Peptic Ulcer Disease

## 8 Declarations

### Ethics approval and consent to participate

The need for Institutional Review Board Approval at our institution (University of Texas Southwestern Medical Center) was waived for this study as all data used for this project had previously been generated as part of either The Cancer Genome Atlas Project or UK Biobank and none of the results reported in this manuscript can be used to identify individual patients.

### Consent for Publication

Not applicable

### Availability of data and materials

TCGA datasets analyzed during the current study are available at the Genomic Data Commons of the National Cancer Institute, https://gdc.cancer.gov/. UK Biobank datasets analyzed during the current study are available from UK Biobank, but restrictions apply to the availability of these data, which were used under license for the current study, and so are not publicly available. Data are however available from the authors upon reasonable request and with permission of UK Biobank. All other datasets used and/or analyzed during the current study are available from the corresponding author on reasonable request.

### Competing Interests

The authors declare that they have no competing interests

### Funding

This work was funded in part by the American Federation for Aging Research (AFAR) and the National Institute on Aging (NIA) through the Medical Student Training in Aging Research (MSTAR) program (GG) and Cancer Prevention and Research Institute of Texas (CPRIT) RR170079 (BL). This work was also supported by the National Institutes of Health [5P30CA142543, 5R01GM126479, 5R01HG008983] and Cancer Prevention & Research Institute of Texas [CPRIT RP190107] (XZ).

### Authors’ Contributions

B.L. proposed the initial idea of using exome data for KIR typing. G.F.G. and B.L. conceived and designed KIRCLE. G.F.G. performed the benchmarking analysis, as well as the association analysis between KIR alleles and molecular and clinical correlates. X.Z. helped with data access and management. G.F.G. and B.L. wrote the manuscript. D.L. and X.Z. offered suggestions and edited the manuscript. B.L. led the project. All authors read and approved the final manuscript.

## Acknowledgements

This research has been conducted using the UK Biobank Resource under Application Number 21237. We are grateful for support from the UK Biobank Access Management Team for their assistance in accessing data from UK Biobank.

## 10 Figures Legends

<u>Supplementary Figure S1. Validation of KIRCLE’s Consistency.</u> Contour plots demonstrating the effect of varying *p* and *t* on consistency of genotype calling, as quantified by entropy, for **(a)** KIR2DL2; **(b)** KIR2DL3, KIR2DL5B, KIR2DS2, & KIR3DS1; **(c)** KIR2DL5A; **(d)** KIR2DS1; **(e)** KIR2DS3; **(f)** KIR2DS4; **(g)** KIR2DS5; **(h)** KIR3DL1; **(i)** KIR3DL2; **(j)** and KIR3DL3.

<u>Supplementary Figure S2. Validation of KIRCLE’s Accuracy.</u>

**(a)** Confusion matrix depicting KIRCLE’s consistency in KIR genotype inference between 531 samples and their biological replicates in TCGA. **(b)** Average entropy versus average depth of coverage in TCGA for each KIR gene. **(c)** Comparison of allele frequency ranks among the 9 KIR3DL2 alleles observed in the US NMDP with their frequency ranks in TCGA (left) and UK Biobank (right).

<u>Supplementary Figure S3. KIR Correlations with Molecular Markers.</u> **(a)** Heatmap of the log2-odds-ratios of KIR allele correlations with HLA alleles in UK Biobank. Correlations with Fisher’s Exact FDR > 0.25 were masked. **(b)** Volcano plot of KIR allele correlations with HLA alleles in TCGA. **(c)** Heatmap of log2-median-fold-changes in tumor immune infiltrate composition estimates stratified by KIR alleles in TCGA. Mann-Whitney-U FDR > 0.25 correlations were masked. **(d)** Volcano plot of KIR allele correlations with differences in tumor immune infiltrate composition in TCGA. **(e)** Heatmap of log2-median-fold-changes in other immune-related molecular signatures and markers stratified by KIR alleles in TCGA. Mann-Whitney-U FDR > 0.25 correlations were masked. **(f)** Volcano plot of KIR allele correlations with differences in other immune-related molecular signatures and markers as measured and characterized in TCGA.

<u>Supplementary Figure S4. KIR Correlations with Other Clinical Variables in UK Biobank.</u> **(a)** Increased likelihood of loose teeth was observed among individuals possessing at least one copy of the KIR3DL3*002 allele compared to those without it. **(b)** Quantitative blood analysis of individuals homozygous for KIR3DL2*010 revealed Increased reticulocyte percentage compared to those without the allele. **(c)** Decreased waist circumference was observed in individuals possessing at least one copy of KIR2DL3*010. **(d)** Lower age of cancer diagnosis was observed among KIR2DS3*002 heterozygotes. **(e)** Lower age of Chronic Obstructive Pulmonary Disease (COPD) diagnosis was observed among KIR3DL2*008 heterozygotes. **(f)** Increased duration of sleep was observed in individuals possessing at least one copy of KIR2DL4*032.

## Notes

### Competing Interest Statement

The authors have declared no competing interest.

