## Supplemental Information for "KIRCLE: An Analysis of Variations in KIR Genes in The Cancer Genome Atlas and UK Biobank"

**11 Supplementary Information**

**Obtaining KIR genomic coordinates**

Using UCSC Table Browser (<https://genome.ucsc.edu/cgi-bin/hgTables>) and the following inputs, we obtained the genomic coordinates of 15 KIR genes in human reference genome GRCh37/hg19:

assembly: “Feb. 2009 (GRCh37/hg19)”

group: “All Tables”

table: “refGene”

identifiers: “KIR2DL1, KIR2DL2, KIR2DL3, KIR2DL4, KIR2DL5A, KIR2DL5B, KIR2DS1, KIR2DS2, KIR2DS3, KIR2DS4, KIR2DS5, KIR3DL1, KIR3DL2, KIR3DL3, KIR3DS1”

**KIRCLE’s Allele Database Creation**

Reference fastas for documented KIR gene variants were obtained from the European Bioinformatics Institute (<https://www.ebi.ac.uk/ipd/kir/genes.html>). Pseudogenes KIR2DP1 and KIR3DP1 were excluded from our final analysis. References were downloaded from the IPD-KIR Database version 2.8.0 in April 2019. 15 total BLAST databases were created for each set of KIR alleles corresponding to one KIR gene.

**Whole Exome Sequencing data**

For analysis of TCGA samples, 10332 WES bam files were downloaded from the Genomic Data Center (GDC) Commons using the GDC’s data transfer tool (<https://gdc.cancer.gov/access-data/gdc-data-transfer-tool>).

For analysis of UK Biobank samples, 49950 WES CRAM files were downloaded using the ukbfetch file transfer tool (<https://biobank.ctsu.ox.ac.uk/crystal/crystal/docs/ukbfetch_instruct.html>).

**Validation of KIR Allele Frequency**

United States National Marrow Donation Program (NMDP) Allele Frequencies were obtained from the Allele Frequency Net Database: <http://www.allelefrequencies.net/>

**Correlation of KIR Alleles with Other Molecular Markers**

Besides associations with HLA alleles, we also assessed for a number of other associations between KIR alleles and various molecular markers previously characterized as part of the PanCanAtlas efforts and demonstrated statistical significance for several of these associations (**Supplementary Figure S2c-f**). Correlates tested include CIBERSORT-estimated proportions of infiltrating immune cell subtypes in the tumors represented in TCGA, as well as gene expression signatures and other molecular markers.[30]

**UK Biobank Correlate Data**

The following data-field IDs were obtained and used for correlation analysis with KIRCLE-generated KIR genotypes:

41202 – ICD10 Primary Diagnosis, Collated from Hospital Inpatient Records

41204 – ICD10 Secondary Diagnosis, Collated from Hospital Inpatient Records

40011 – Tumor Histology, Provided by the Cancer Registry

6149 – Self-Reported Dental/Oral Problems

2178 – Self-Reported Overall Health

2188 – Self-Reported Long-standing Illness, Disability, or Infirmity Status

40012 – Tumor Behavior, Provided by the Cancer Registry

6154 – Self-reported use of medications for either pain, constipation, or heartburn relief

1160 – Self-Reported Sleep Duration

137 – Self-Reported Number of Medications Being Taken

40008 – Cancer Diagnosis Age, Provided by the Cancer Registry

134 – Self-Reported Number of Cancerous Conditions/Diagnoses

135 – Self-Reported Number of Non-Cancerous Conditions/Diagnoses

21001 – Measured Body Mass Index

12144 – Measured Height

48 – Measured Waist Circumference

30000 – 30300 – Quantitative Blood Analysis From Obtained Blood Sample

22150 – Self-Reported Age of COPD Diagnosis

22152 – Self-Reported Age of alpha-1 antitrypsin deficiency diagnosis

22159 – Self-Reported Age of asbestosis diagnosis

22147 – Self-Reported Age of asthma diagnosis

22154 – Self-Reported Age of bronchiectasis diagnosis

22149 – Self-Reported Age of chronic bronchitis diagnosis

3761 – Self-Reported Age of hayfever, rhinitis, or eczema diagnosis

**Correlation Analyses**

For categorical variables, such as HLA-I status, we used Fisher’s Exact tests to perform correlations with KIR allele status.

For continuous variables, such as age of onset of allergic reaction, we used 1-way ANOVA tests and Mann-Whitney-U tests to perform correlations with KIR allele status.

False-Discovery-Rate correction was performed using the Bonferroni method.

For correlations with HLA alleles in UK Biobank, alleles with a probability estimate > 0.75 were considered to have one copy of the HLA-I allele, whereas, those with a probability estimate >1.5 were considered to have 2 copies of the HLA-I allele.

For analyses where we considered only high-frequency alleles, we excluded all alleles with a population frequency <1%, as determined by counting the total occurrences of the allele and dividing by two-times the number of individuals in the dataset being considered.

For correlations with ICD10 codes in UK Biobank, all ICD10 primary and secondary diagnoses (datafields 41202 and 41204) were considered. Additionally, we excluded the following blocks and, thus, any ICD10 codes within them: A0-A99, B0-B99, F0-F99, H0-H99, O0-O99, P0-P99, Q0-Q99, R0-R99, S0-S99, T0-T99, U0-U99, V0-V99, W0-W99, X0-X99, Y0-Y99, Z0-Z99, D50-D53, E40-E46, E50-E64, J00-J06, J09-J18, J20-J22, J60-J70, L00-L08, L55-L59, M00-M03, and N20-N23.

**Software Used for Computational Analysis**

KIRCLE was run using samtools 1.3.1 and Nucleotide BLAST v2.8.1+.

All other computational analyses were performed using Python v3.6.10 augmented with the following packages: numpy v1.18.1, pandas v1.0.1, scipy v1.4.1, sklearn v0.22.1, pysam v0.9.1, matplotlib v3.2.1, and seaborn v0.10.0.

Scripts were run on Red Hat Enterprise Linux Server release 7.4 (Maipo) on the BioHPC-Nucleus Supercomputer at UT Southwestern Medical Center.
