## Supplementary figures and images for "KIRCLE: An Analysis of Variations in KIR Genes in The Cancer Genome Atlas and UK Biobank"

### Supplemental Figures

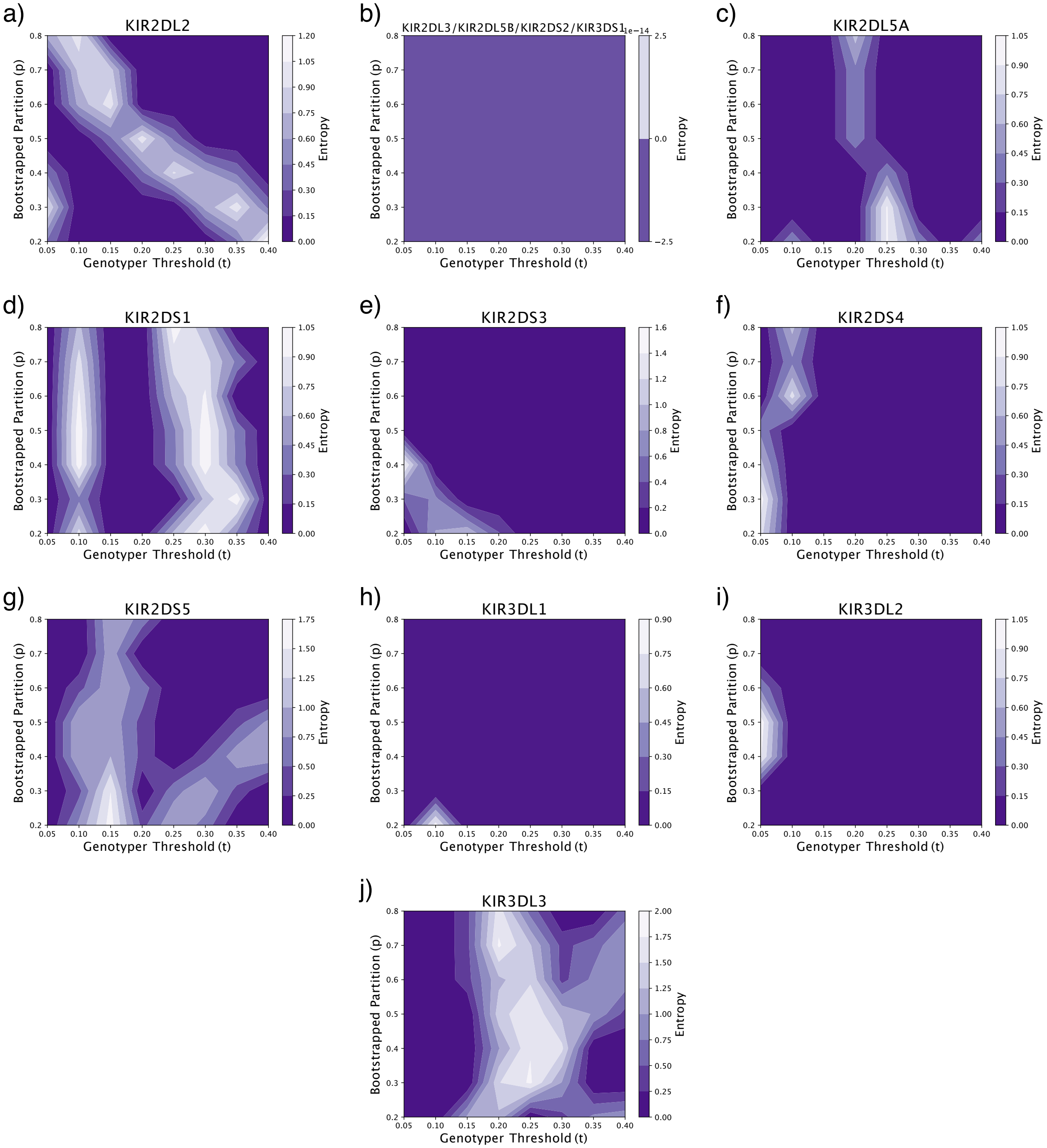

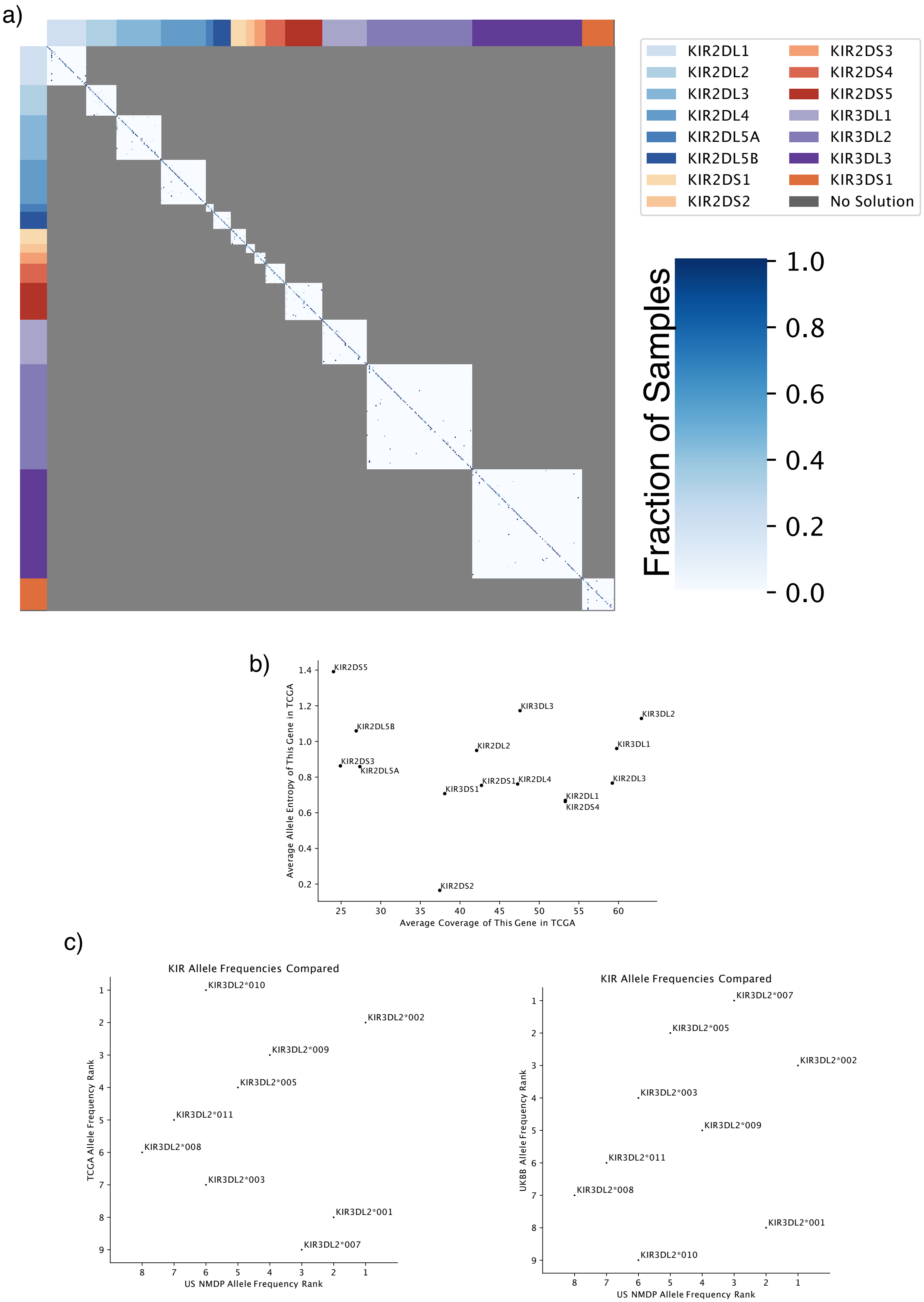

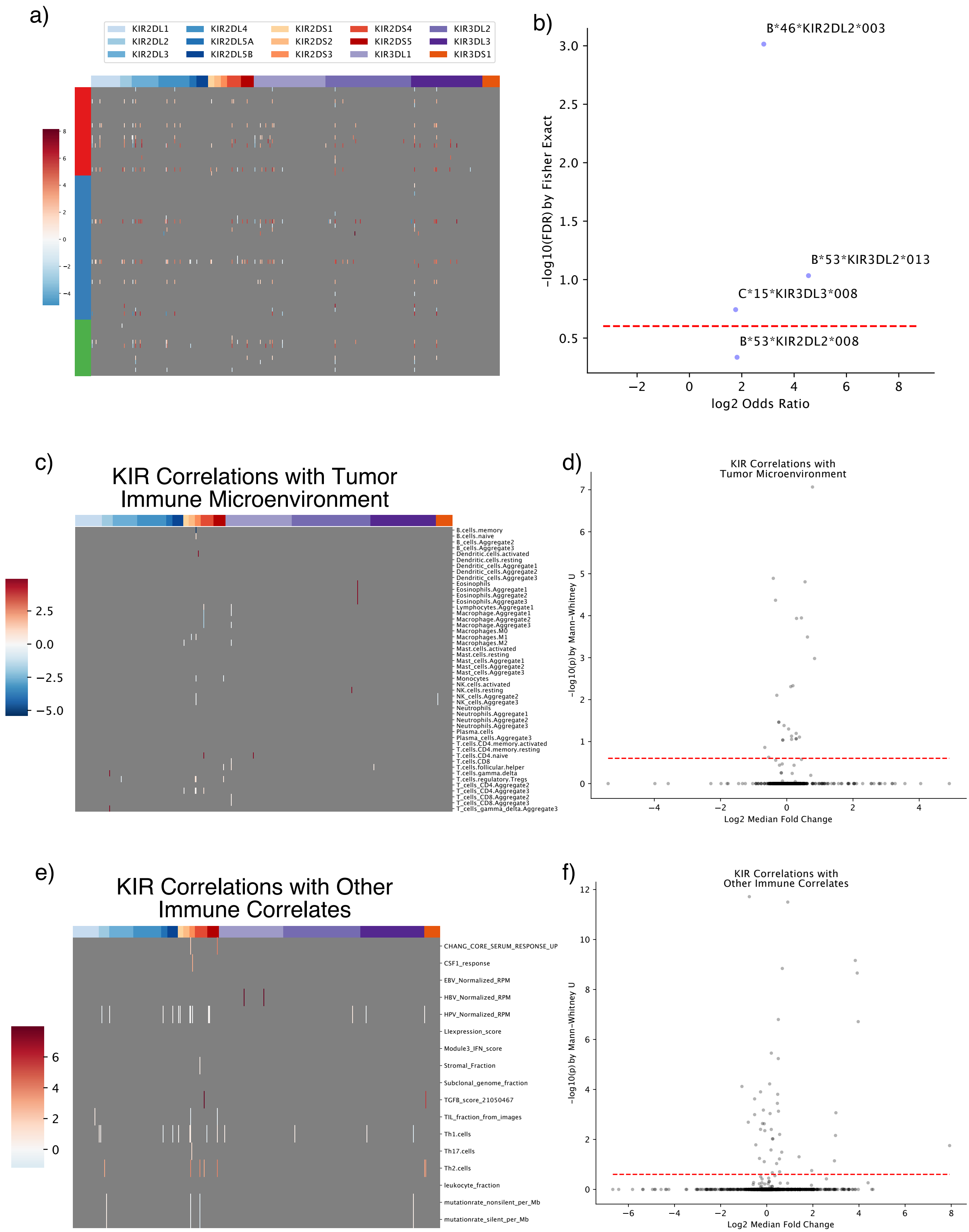

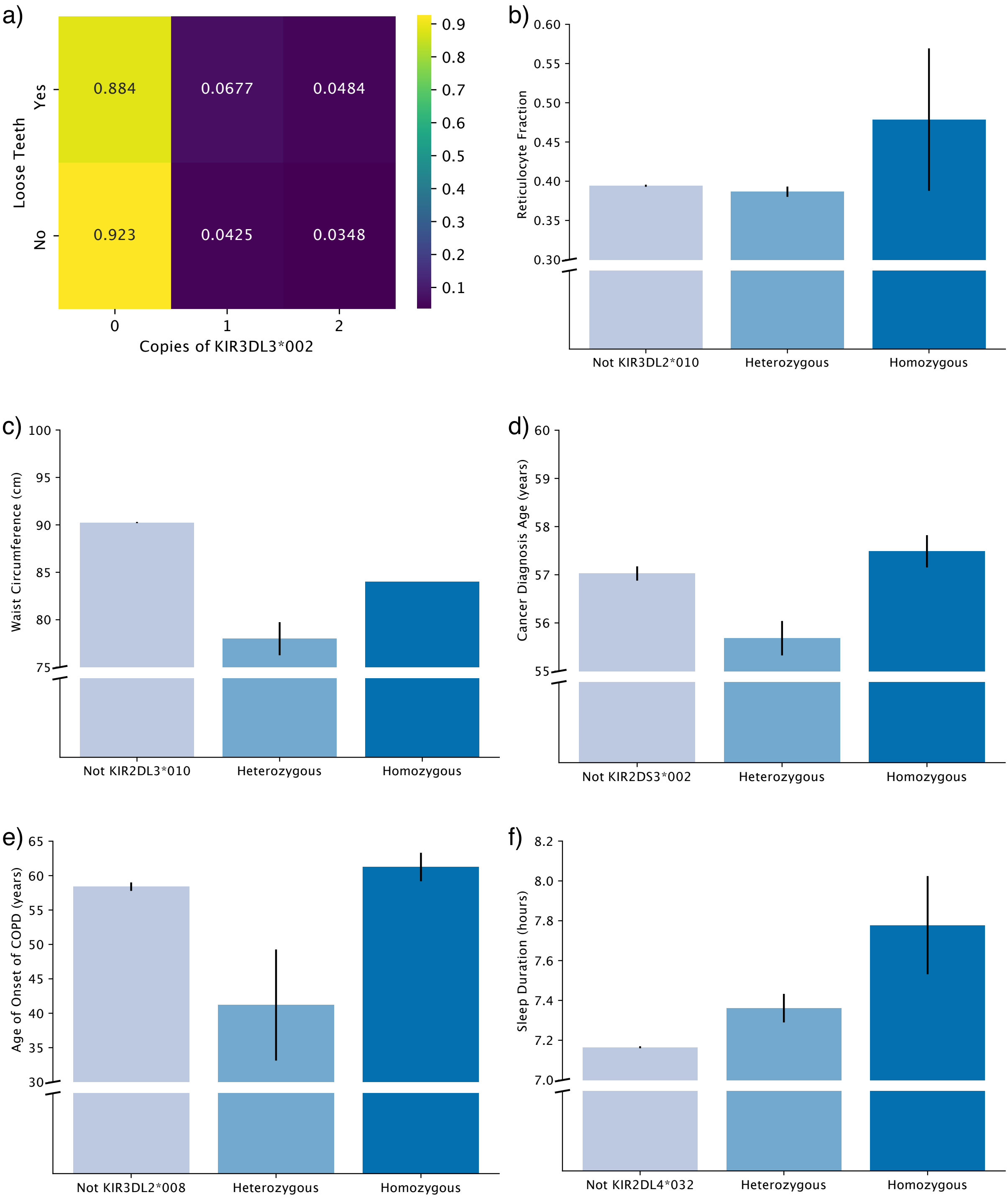
